# *In vivo* cloning of β 1-4 endoglucanase gene of *Serratia liquefaciens* using Muduction and its *in silico* analysis

**DOI:** 10.1101/283226

**Authors:** S. Gowri Sankar, J. Asnet Mary, S. John Vennison, A. Alwin Prem Anand

**Affiliations:** Department of Biotechnology, Anna University - BIT Campus, Tiruchirappalli – 620024, Tamil Nadu, India.; Research Center for Biological Sciences, Naesam Trust, Ellis Nagar, Madurai – 625016, Tamil Nadu, India.; Department of Molecular Biology and Diagnostics, Centre for Research in Medical Entomology (CRME), Madurai – 625002, Tamil Nadu, India; Department of Zoology, Fatima College, Madurai – 625018, Tamil Nadu, India.

**Keywords:** bacteriophages, biotechnology, biodegradation, enzymes, enzymology

## Abstract

Cellulose is the major structural component in the plant cell wall. The bio-degradation of cellulose molecules is facilitated by cellulase. In the present study, *in vivo* cloning of cellulase (β-1, 4 endoglucanase) gene from a cellulolytic bacterium *Serratia liquefaciens* into *E. coli* DH5α has been performed using mini-Mu phage transduction. The enzyme activity of cloned endoglucanase was 81.2U/mg at optimum temperature (40°C) and 80.2U/mg at optimum pH7, while the wildtype has 65.9U/mg and 64.9U/mg respectively. The conserved domain analysis shows that *S. liquefaciens* endoglucanase belongs to GH8 family. The nucleotide sequence analysis of wildtype and cloned endoglucanase shows that mutations were found at residues 51(Lys - Asn), 203(Trp-Cys), 246(Thr-Iso), 260(Gly-Ala) and 288(Phe-Leu). The structural analysis shows the active site of wildtype endoglucanase is a narrow groove which lies parallel to the central axis, whereas cloned endoglucanase is broad and tilted to ∼70° from the central axis. The increased enzyme activity in the cloned endoglucanase is due to the structural modification conferred by changes in amino acid resulting in widening of the cleft in the active site.

## Introduction

Cellulose is the most abundant material on earth, which can be harnessed as sustainable source of biofuel. Cellulose is a biopolymer made of β 1,4 glycosidic linkages of glucose. Endoglucanases (EC 3.2.1.4) are enzymes that cut randomly at the internal amorphous in the cellulose polysaccharide chain (Lynd et al. 2002; Juturu and Wu 2014). The endoglucanase are produced by bacteria, fungi (Doi and Kosugi 2004) and in insects, where some of them are endogenous origin (Watanabe et al. 1998; Watanabe and Tokuda 2001) and some are in symbiotic relations with microbes (Watanabe and Tokuda 2010). From our previous experiment with *Bombyx mori*, bacteria capable of degrading cellulose were isolated from the digestive tract (Alwin Prem Anand et al. 2010). Among the isolates, *Serratia liquefaciens* has been chosen for cloning of endoglucanase gene using mini-Mu system; a bacteriophage based *in vivo* cloning.

Gene cloning using mini-Mu replicon is a simple and efficient *in vivo* approach for prokaryotic gene cloning (Groisman and Casadaban 1986; Stojiljkovic et al. 1994). Bacteriophage Mu and related mutator phages are the largest and most efficient known transposable elements. During the replication process it transposes to more than fifty different sites in the genome within an hour (Taylor 1963; Howe and Bade 1975; Bukhari 1976; Groisman and Casadaban 1987b; Groisman and Casadaban 1987a) and frequently cause mutations of the host bacteria (Taylor 1963). High frequency of transposition and DNA packaging property make Mu phage ideal for several genetic procedures including cloning (Akhverdyan et al. 2011; Harshey 2014; Pulkkinen et al. 2016a; Pulkkinen et al. 2016b). The mini-Mu phage has been used: for target sequence specific insertional mutagenesis of the HSV genome (Jenkins et al. 1985), to clone nitrofuran reductase gene into *E. coli* (Kumar and Jayaraman 1991) and, to identify the regions encoding the structure and regulatory protein gene of tetracycline resistant determinants of *Proteus mirabilis* (Magalhaes and Castilho 1997). As Mu based system holds several advantages over other *in vitro* manipulation technologies, we are interested in using the mini-Mu system for *in vivo* cloning of β-1,4 - endoglucanase (cellulase) gene from cellulolytic bacteria.

The cloning of endoglucanase gene from *S. liquefaciencs* to *E. coli* using muduction was successful. The experimental evidence shows that the enzyme activity of cloned endoglucanase has been increased up to 24% when compared to endoglucanase of wildtype *S. liquefaciens*. Both endoglucanases were analyzed by *in silico* methods to reason out (unravel) the enhanced enzyme activity of cloned endoglucanase. Since the cloning was done using Mu phage, it was anticipated that some mutation (Taylor 1963; Mizuuchi and Craigie 1986) would have been introduced in the endoglucanase gene, which might change the sequence and structure of endoglucanase so as to increase the interactions between substrate and active site. In order to understand the role of mutated amino acids, the protein sequences were compared, theoretical models of the wildtype and cloned endoglucanase were generated and the structural properties were analyzed.

## Materials and Methods

### 1. Strains and plasmids

The source strain of β-1,4-endoglucanase gene is the cellulolytic bacterium *S. liquefaciens*. Originally, this strain was isolated from the digestive tract of *Bombyx mori* during the fifth instar larval stage. The isolation and characterization has been discussed earlier (Alwin Prem Anand et al. 2010) and further confirmed by 16S rRNA sequencing (*Genebank: KT253149*). The cloning and expression was done using *E. coli* DH5α. *E. coli* AB1157 Mu cts and strain carrying plasmid pHYD803 were kind gift from Prof. Munavar, Madurai Kamaraj University, India. Strains (Table 1) were maintained in appropriate antibiotic supplemented media until used.

**Table 1.** Strains and plasmids used. This table shows the strains and plasmid used in the experiment.

| Strains/plasmids | Properties | Reference |
| --- | --- | --- |
| *E. coli* AB1157<br>(harbouring Mu<br>*cTs*) | *thr-1 ara-14 leuB6 Δ (gpt-proA) 62*<br>*lacY1 tsx-33 supE44 galK2 hisG4*<br>*rpoS338 rpsL31 kdgK51 xyl-5 mtl-1*<br>*argE3 thi-1* (Mu *cts*) | Saroja and Gowrishankar<br>(1996) |
| pHYD803 | Carries Mini-Mu phage and Kan<sup>r</sup><br>selection marker | Saroja and Gowrishankar<br>(1996) |
| *E. coli* DH5α | F– φ80lacZZ ΔlacZYA-argF) U196<br>endA1 recA1 hsdR17 (rk–, mk+)<br>supE44 thi-1 gyrA96 relA1 phoA |  |

### 2. Chemicals, media and general laboratory techniques

Lysogenic Broth (LB) was obtained from Himedia Laboratories. All other chemicals such as CMC, Congo red, MgSO_4_, CaCl_2_, Chloroform, and NaCl were obtained from Merck Laboratories. Composition of minimal media (Berg’s Agar) containing carboxyl methylcellulose was described elsewhere (Berg et al. 1972). All other chemicals and reagents used were of molecular grade. General laboratory techniques were carried out following Maniatis and colleagues (1982).

### 3. *In vivo* gene cloning

#### Phage lysate preparation and Mu lysogen construction

Mu cts lysates were prepared using *E. coli* AB1157 [Mu*c*(Ts)]/pHYD803 by method described earlier with some minor modifications (Groisman and Casadaban 1986). Early exponential phase *E. coli* cells harboring phages grown at 28°C were shifted to 42°C until lysis occurs. Chloroform (10% vol/vol) was added and the cell pellets were removed by centrifugation. MgSO_4_ and CaCl_2_ to a final concentration of 2mM and 0.2mM were added to the lysate. Mu lysogens were constructed by spotting fresh Mu cts lysate on bacterial lawn (*Serratia liquefaciens*) and kept overnight at 30°C. The lysate was collected as described above, for muduction in recipient cells (*E. coli* DH5α).

#### Mu-duction

The method described by Groisman and Casadaban (1986) was followed. For infection, 100µl of overnight culture of recipient cells - *E. coli* DH5α cells, were mixed with equal volumes of phage and kept for 30 min at 30°C for phage adsorption, without shaking. Then 2 ml of fresh LB broth was added and the culture was further shaken at 30°C for 1.5 hr to allow phenotypic expression. The cells were washed twice with normal saline before plating in minimal media containing CMC and Kanamycin. The cloned *E. coli* strain degrading CMC with high efficiency was selected and used in further experiments.

### 4. Enzyme (1,4-beta endoglucanase) assay

#### Screening of Clones with 1,4-beta endoglucanase activity

The screening for clones were done using Congo red overlay method (Teather and Wood 1982). Plates were flooded with 0.1% aqueous congo red for 10 minutes and washed with 1M NaCl solution for two times. The clearing zone around the colony indicates the hydrolysis of CMC (Wood 1980; Ruijssenaars and Hartmans 2001).

#### Quantitative analysis of 1,4-beta endoglucanase activity (DNS method)

The quantitative analysis of endoglucanase activity was assayed by measuring the amount of reducing sugar liberated from assay mixture containing 1% CMC in 0.05 M sodium acetate buffer (pH 7.0). The reaction was arrested by DNS and the absorbance was measured at 540 nm (Miller 1959).

#### Enzyme production and purification

Enzyme production was done in 500ml of Erlenmeyer flask containing 100ml of medium supplemented with 1% CMC and inoculated with 10^5^ cells/ml and incubated with shaking for 36hrs at the different temperatures. Aliquotes were withdrawn for every 4 hr and enzyme activity was measured by DNS method. After incubation, the culture was centrifuged at 14,000 × g for 15 min at 4°C and the supernatant was used as crude enzyme.

Enzyme purification was done by ammonium sulfate purification method. Briefly ammonium sulfate was added with culture supernatant to attin 60% (W/V) and kept for 12 hrs at 4°C. The mixture was centrifuged at 10,000 × g and the precipitate was dissolved in 50mM-Tris HCl and dialyzed against the same solution. The precipitate was loaded in Sephadex G-50 coloumn and the elution was carried out using linear gradient of 0-1M NaCl. The fractions were screened for endoglucanase activity and pooled.

#### SDS-PAGE and Zymogram analysis for 1,4-**β** endoglucanase

The samples were boiled for 5 minutes with Laemmli sample buffer with ß Mercaptoethanol at a 1:1 ratio. Samples were electrophoretically separated on an SDS PAGE gel Gels were fixed for 45 mins in 30% methanol, 10% acetic acid and then stained with 0.006% Coomassie Brilliant Blue G-250, 10% acetic acid for two hours. Destaining was performed in 10% acetic acid for 90 mins.

Zymogram was done in 10% polyacrylamide gel containing 1% CMC (native PAGE). The gel was refolded in 15% isopropanol in phosphate buffered saline followed with two washes and stained with 0.1% Congo red for 1 hr and subsequently washed with 1M NaCl solution.

#### Sequencing of beta-1, 4-endoglucanase gene of *Serratia* (wildtype) and Mini Mu phagemid (cloned)

For sequencing the cloned beta 1-4 endoglucanase gene, primers (MLF-CTGGGCGGATAGCCTGATTA and MRR-TCGCGTTCATGGTAATTTCA) were designed based on pHYD803 sequences. The genomic DNA of S. *liquefaciens* was used as a template and primers (fwd 5’-ATGCCTGCGGCATTGGCCTCTG-3’; rev 5’-TCACGCATTGGCCGCCCCAGGA-3’) were designed based on β 1-4 endoglucanase sequences deduced from pHYD803. The DNA sequencing was done using Sequencing technology – 454 commercially. Sequences were aligned using Chromas (v.2.4.1) software and submitted to NCBI.

### 5. *In silico* analysis

#### Sequence analysis

Sequence alignment of the nucleotide sequences of endoglucanase from *S. liquefaciens* (*GenBank: KF032300.1*) and cloned endoglucanase (Mini mu Phagemid – *GenBank: KF032301.1*) were performed with EMBOSS Water to find one or more alignments describing the most similar regions within the sequences. EMBOSS Water uses the Smith-Waterman algorithm to calculate the local alignment of a sequence to one or more other sequences (McWilliam et al. 2013). Pair-wise alignment options were set as default. Phylogenetic relationship of cellulase proteins among *Serratia* sp. was predicted using MEGA6.0 (Tamura et al. 2013). The bootstrap consensus tree inferred from 500 replicates has been taken to represent the evolutionary history of all species. The evolutionary distances were estimated using the Neighbor-Joining method with Kimura 2-parameter model.

#### Structural comparison of Endoglucanase of *Serratia* and Mu phage

To associate the sequence and functional relationship between the wildtype endoglucanase (*S. liquefaciens*) and cloned endoglucanase (Mu Phagemid), the sequences were compared for their physico-chemical properties such as molecular weight, isoelectric point, amino acid composition, hydropathicity and the number of disulphide bridges. The disulphide bonds were predicted using DiANNA and CYSPRED tools. DiANNA predicts the disulfide connectivity by running PSIPRED and then PSIBLAST (Fariselli et al. 1999; Ferre and Clote 2005).

The secondary structures of endoglucanase protein of *Serratia* and Mini Mu Phagemid were predicted using PSIPRED (McGuffin et al. 2000). The prediction of three-dimensional structures of endoglucanase proteins was performed using I-TASSER. The parameters were set as default with no restraints on selection of templates (Roy et al. 2010). The stereochemical qualities of the predicted structures were analyzed by PROCHECK. Both the predicted structures were compared with DaliLite server available at EBI (Holm and Park 2000), which is a pairwise structure comparison program. The predicted structures were analyzed for the number of hydrogen bonds and molecular surface area. It was detected by VADAR (Version 1.1) which computes volume, area, and dihedral angles in the protein structures from their PDB coordinate data (Willard et al. 2003).

#### Results

##### Wildtype and cloned endoglucanase of *S. liquefaciens*

The cellulolytic bacterium *S. liquefaciens* used here was isolated from the digestive tract of fifth instar larval stage of *B. mori* (Alwin Prem Anand et al. 2010). Prior to cloning of the endoglucanase gene from *S. liquefaciens* using muduction, the *E. coli* DH5α was checked on CMC plate for any endoglucanase activity and were found to be none. *S. liquefaciens* has the clearing zone of about 6 mm in CMC medium. After muduction, we have obtained 44 *E. coli* isolates with cellulolytic activity. Among the 44 isolates, four clones (Clone #1 = 6 mm, Clone #2 = 7 mm, Clone #3 = 7 mm and Clone #4 = 10 mm) have increased enzyme activity than the wildtype and the remaining clones have reduced enzyme activity compared to the wildtype i.e., less than 6 mm. Among the four clones, the clone (Clone #4) with highest endoglucanase activity has been used for further experiments.

Both the cloned endoglucanase (*E. coli* Mu phagemid) and wildtype (*S. liquefaciens*) endoglucanase has a molecular mass of about 32kDa (Fig. 1A). In both cases, the optimum temperature is 40°C (Fig. 1B) and the enzyme activity of wild type was found to be 65.9 U/mg and the cloned gene was 81.2 U/mg, which is approximately 24% higher than the wild type. Both cloned and wildtype endoglucanase has an optimum pH about 7 with an enzyme activity of 64.9U/mg in wildtype and 80.2U/mg in cloned endoglucanase (Fig.1C).

**Fig. 1.**
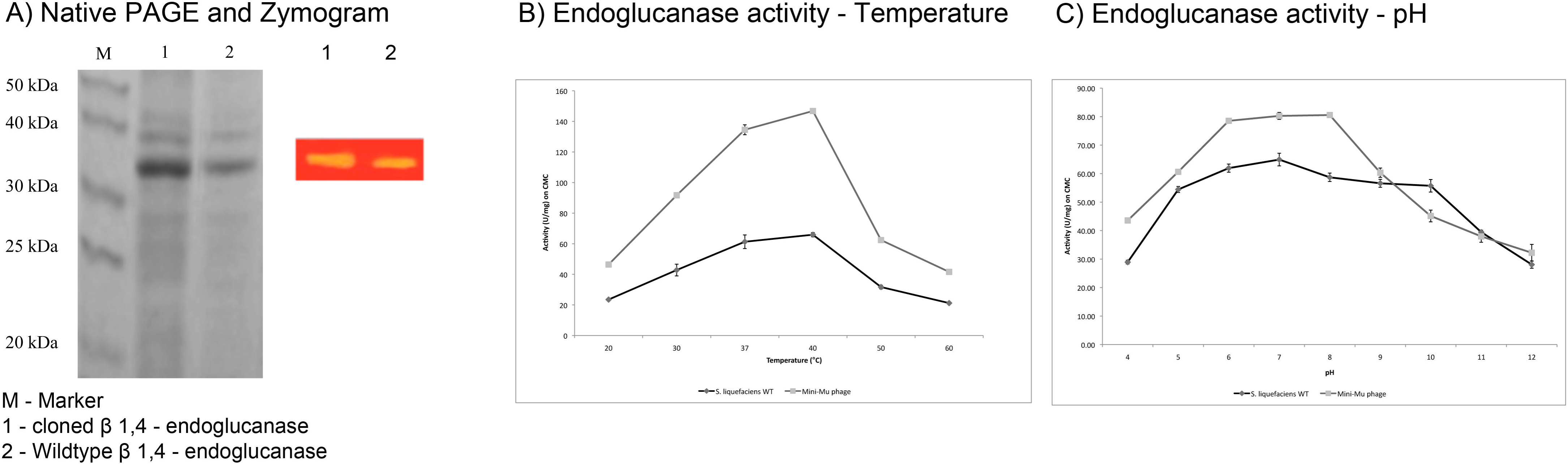
Functional aspect of wildtype (*S. liquefaciens*) and cloned (Mu phagemid) endoglucanase. A) The native PAGE showing the molecular mass of the cloned and wildtype endoglucanase as 32kDa. The coloured picture (zymogram) shows the enzyme activity of cloned and wildtype endoglucanase. B) The endoglucanse activity of the wildtype (*S. liquefaciens*) and cloned endoglucanase (mini-Mu phagemid) shows there is a 24% increase in cloned endoglucanase. Both the endoglucanases are active at 37-40°C. C) The endogulcanase activity of the wildtype (*S. liquefaciens*) and cloned endoglucanase (mini-Mu phagemid) shows there is a 24% increase in cloned endoglucanase. Both the endoglucanases are active at pH6-8. One enzyme unit (U) was defined as the enzyme amount, which releases 1μM of glucose equivalent from substrate per minute.

In order to understand the increased activity of the cloned endoglucanase, we have sequenced and done computational analysis by comparing the cloned endoglucanase and the wildtype of *S. liquefaciens.* The nucleotide sequences of endoglucanase from *S. liquefaciens* (*GenBank: KF032300.1*) and cloned endoglucanase (Mini mu Phagemid – *GenBank: KF032301.1*) are available in Genbank.

##### Sequence analysis of wildtype and cloned endoglucanase

The sequence analysis by conserved domain database (CDD) (Marchler-Bauer et al. 2015) shows that the conserved domain of *S. liquefaciens* endoglucanase belongs to glycosyl hydrolyses family 8 (GH8, Fig. 2A). The nucleotide sequence analysis of endoglucanase of wildtype and the cloned gene revealed five synonymous and three non-synonymous mutations. The mutations were found at residues 51 (Lys - Asn), 203 (Trp-Cys), 246 (Thr-Iso), 260 (Gly-Ala) and 288 (Phe-Leu) (Fig. 2B). We found increased number of amino acids like Alanine, Asparagine, Cysteine, Isoleucine, Leucine and reduced number of amino acids like Glycine, Lysine, Phenylalanine, Threonine and Tryptophan in cloned endoglucanase (Table 2). The phylogenetic relationship of cellulase protein of *Serratia* with other organisms from *Enterobacteriaceae* shows that *S. liquifaciens* distantly related with other organisms whereas *Enterobacter* sp., *Salmonella*, *Pectobacterium*, and *Yersinia* are clustered as one group (Supplementary Figure 1).

**Fig. 2.**
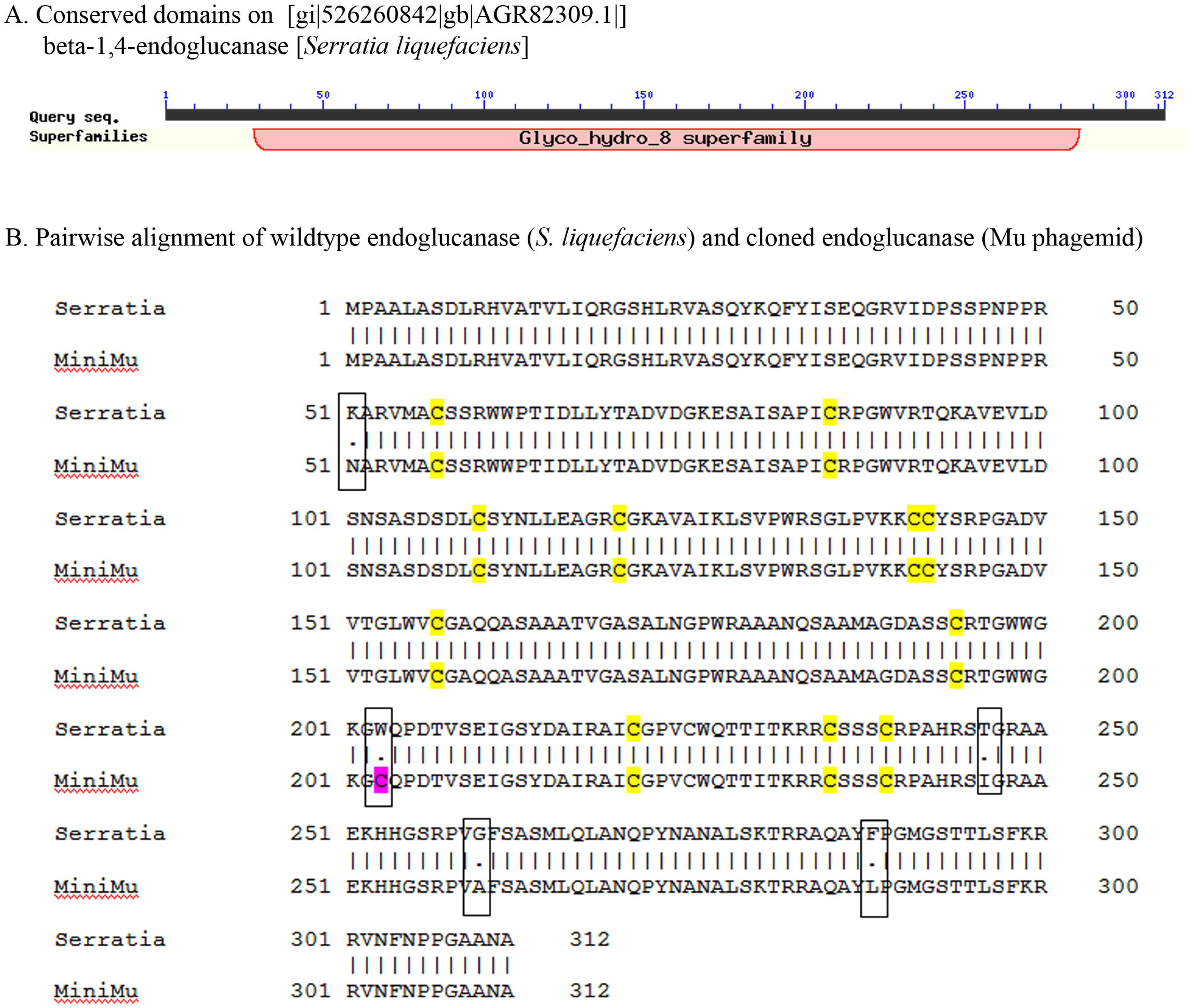
Sequence analysis of wildtype and cloned endoglucanase. **A. This shows the conserved domain on wildtype endoglucanase (*S. liquefaciens*).** The conserved domain has sequence similarities with Glycosyl hydrolyses family 8. **B. Pairwise alignment of wildtype endoglucanase (*S. liquefaciens*) and cloned endoglucanase (Mu phagemid).** The length of both the sequences are same. The sequences differ at five sites, 51^st^ (Lys - Asn), 203^rd^ (Trp-Cys), 246^th^ (Thr-Iso), 260^th^ (Gly-Ala) and 288^th^ (Phe-Leu), which are represented in rectangular boxes. The cysteine residues in wildtype and cloned endoglucanase are highlighted in yellows. In cloned endoglucanase, cysteine is introduced in place of Tryptophan at 203^rd^ position, which is higlighted in pink.

**Table 2.** The Physico-chemical properties of wildtype endoglucanase (S. liquefaciens) and cloned endoglucanase (Mu phagemid). The number of amino acids in both are similar which differ slightly in their molecular weight and isolelectric point. The amino acids differ in the sequences alone are tabulated which comes to maximum of 0.3%. The hydropathicity is reduced to some extent in cloned endoglucanase.

| Properties | Wildtype endoglucanase<br>(*S. liquefaciens*) | Cloned endoglucanase<br>(Mu Phagemid) |
| --- | --- | --- |
| Number of amino acids | 312 | 312 |
| Molecular weight | 33415.9/33417 | 33310.8/33312 |
| Theoretical pI | 9.82 | 9.72 |
| Amino acid composition | Ala (A) 44 14.1%<br>Asn (N) 11 3.5%<br>Cys (C) 12 3.8%<br>Gly (G) 24 7.7%<br>Ile (I) 11<br>3.5%<br>Leu (L) 18 5.8%<br>Lys (K) 13 4.2%<br>Phe (F) 5 1.6%<br>Thr (T) 15 4.8%<br>Trp (W) 10 3.2% | Ala (A) 45 14.4%<br>Asn (N) 12 3.8%<br>Cys (C) 13 4.2%<br>Gly (G) 23 7.4%<br>Ile (I) 12<br>3.8%<br>Leu (L) 19 6.1%<br>Lys (K) 12 3.8%<br>Phe (F) 4 1.3%<br>Thr (T) 14 4.5%<br>Trp (W) 9 2.9% |
| Total number of negatively charged residues (Asp + Glu) | 18 | 18 |
| Total number of positively charged residues (Arg + Lys) | 38 | 37 |
| Grand average of hydropathicity (GRAVY): | -0.292 | -0.253 |

In order to understand whether Cys203 is involved in disulphide bridge formation, prediction was made with DiANNA and CYSPRED, which are disulphide bond prediction tools. In case of wildtype endoglucanase, there is a disulphide bond between Cys120 and Cys142 whereas in cloned endoglucanase, Cys203 the mutated residue is bonded with Cys120 (Table 3). The connectivity of cysteine residues were altered between wildtype endoglucanase (Cys120 – Cys142, Cys141 – Cys235, Cys221 – Cys225) and cloned endoglucanase (Cys120 – Cys203, Cys142 – Cys221, Cys225 – Cys235).

**Table 3.** Disulphide bond Prediction in wildtype endoglucanase (*S. liquefaciens*) and cloned endoglucanase (Mu phagemid). It is predicted by DiANNA and CYSPRED tools. The number of cysteine residues in wildtype endoglucanase and cloned endoglucanase are 12 and 13 respectively. DiANNA prediction shows six disulphide bonds in both wildtype and cloned endoglucanase; but show differences in their connectivity. In cloned endoglucanase, the Trp203 is substituted with Cys203 residue and thereby introducing one new cysteine residue. CYSPRED predicts eight sites as bonding state and three sites as non-bonding state in wildtype endoglucanase whereas in cloned endoglucanase, nine sites have been predicted as bonding state which includes the Cys203, the mutated residue, and 4 sites as non-bonding state.

| Endoglucanase Protein | DiANNA Predicted connectivity in Parentheses | CYSPRED |  |  |
| --- | --- | --- | --- | --- |
|  |  | Cysteine residue with number | Prediction | Reliability |
| *S. liquefaciens* (Wildtype) | 57 - 239 | CYS 57 | NON-Bonding State | 8 |
|  | 85 - 157 | CYS 85 | Bonding State | 3 |
|  | 110 - 194 | CYS 110 | Bonding State | 8 |
|  | 120 - 142 | CYS 120 | NON-Bonding State | 2 |
|  | 141 - 235 | CYS 141 | Bonding State | 2 |
|  | 221 – 225 | CYS 142 | Bonding State | 2 |
|  | (1-12, 2-7, 3-8, 4-6, 5-11, 9-10) | CYS 157 | Bonding State | 5 |
|  |  | CYS 194 | Bonding State | 1 |
|  |  | CYS 221 | Bonding State | 2 |
|  |  | CYS 225 | NON-Bonding State | 6 |
|  |  | CYS 235 | Bonding State | 8 |
|  |  | CYS 239 | Bonding State | 7 |
| Mu Phagemid (Clone) | 57 - 239 | CYS 57 | NON-Bonding State | 8 |
|  | 85 - 157 | CYS 85 | Bonding State | 3 |
|  | 110 - 194 | CYS 110 | Bonding State | 8 |
|  | 120 - 203 | CYS 120 | NON-Bonding State | 2 |
|  | 142 - 221 | CYS 141 | Bonding State | 2 |
|  | 225 - 235 | CYS 142 | Bonding State | 2 |
|  | (1-13, 2-7, 3-8, 4-9, | CYS 157 | NON-Bonding State | 5 |
|  |  | CYS 194 | Bonding State | 1 |
|  | 6-10, 11-12) | **CYS 203** | Bonding State | 8 |
|  |  | CYS 221 | Bonding State | 2 |
|  |  | CYS 225 | NON-Bonding State | 6 |
|  |  | CYS 235 | Bonding State | 8 |
|  |  | CYS 239 | Bonding State | 7 |

##### Structural analysis of cloned and wildtype endoglucanase

To interrogate the structural consequences of amino acid substitutions in their secondary and tertiary structures, the theoretical models of wild and mutant type endoglucanase were generated. The secondary structures for both wildtype and mutated proteins were predicted using PSIPRED (Fig. 3) and the three-dimensional (3D) structure were predicted using I-TASSER (Fig. 4). It was found that the secondary structures of wildtype (Fig. 3A) and cloned endoglucanase (Fig. 3B) remain unaltered.

**Fig. 3.**
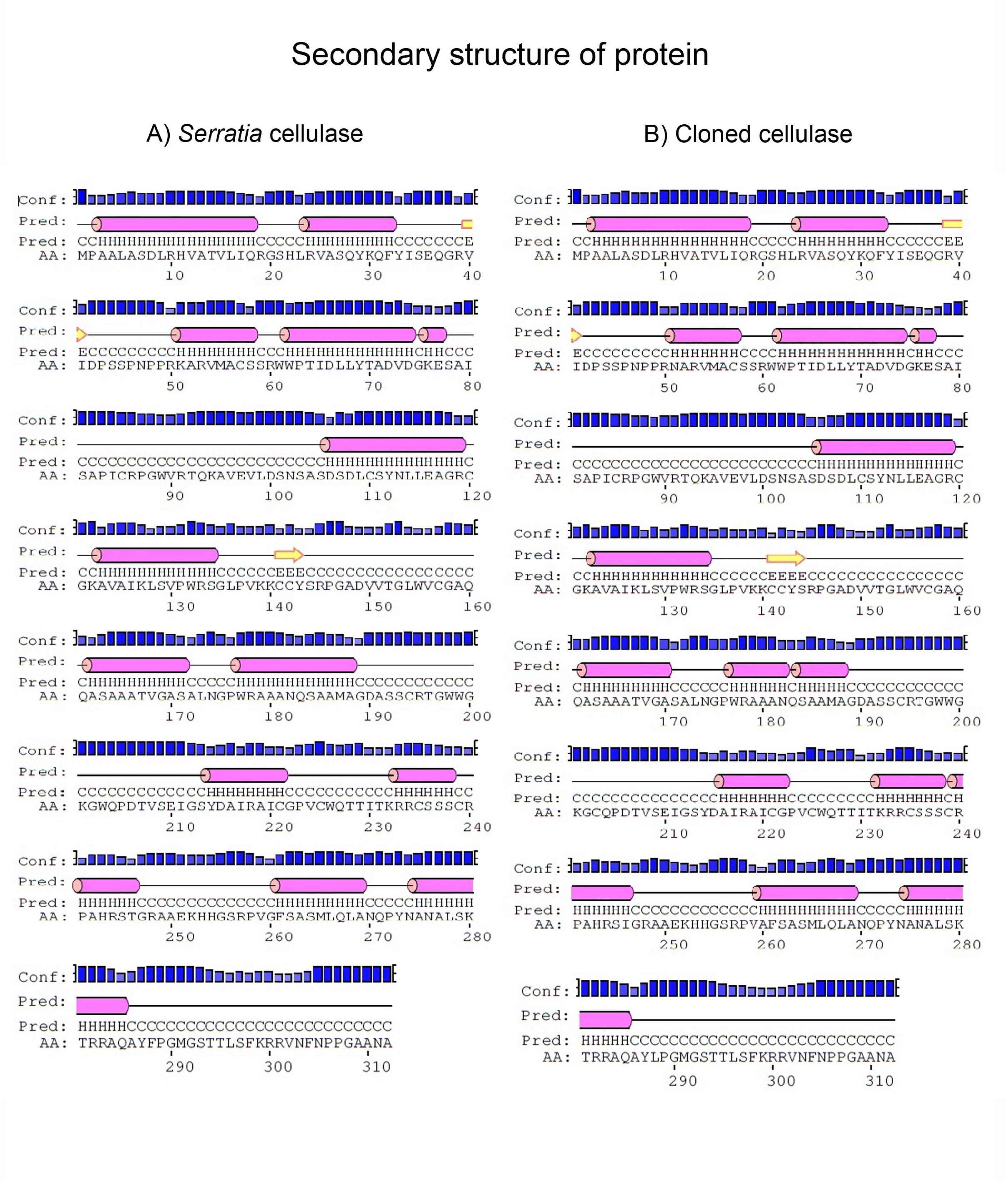
Prediction of Secondary structure of Endoglucanase Protein. A. Endoglucanase of *Serratia* B. Mu Phage. The α-helices are represented as bar in red, beta-sheets in yellow arrow mark. Black horizontal line illustrates loop regions. No significant difference in protein secondary structures of *S. liquefaciens* and Mu Phagemid.

**Fig. 4.**
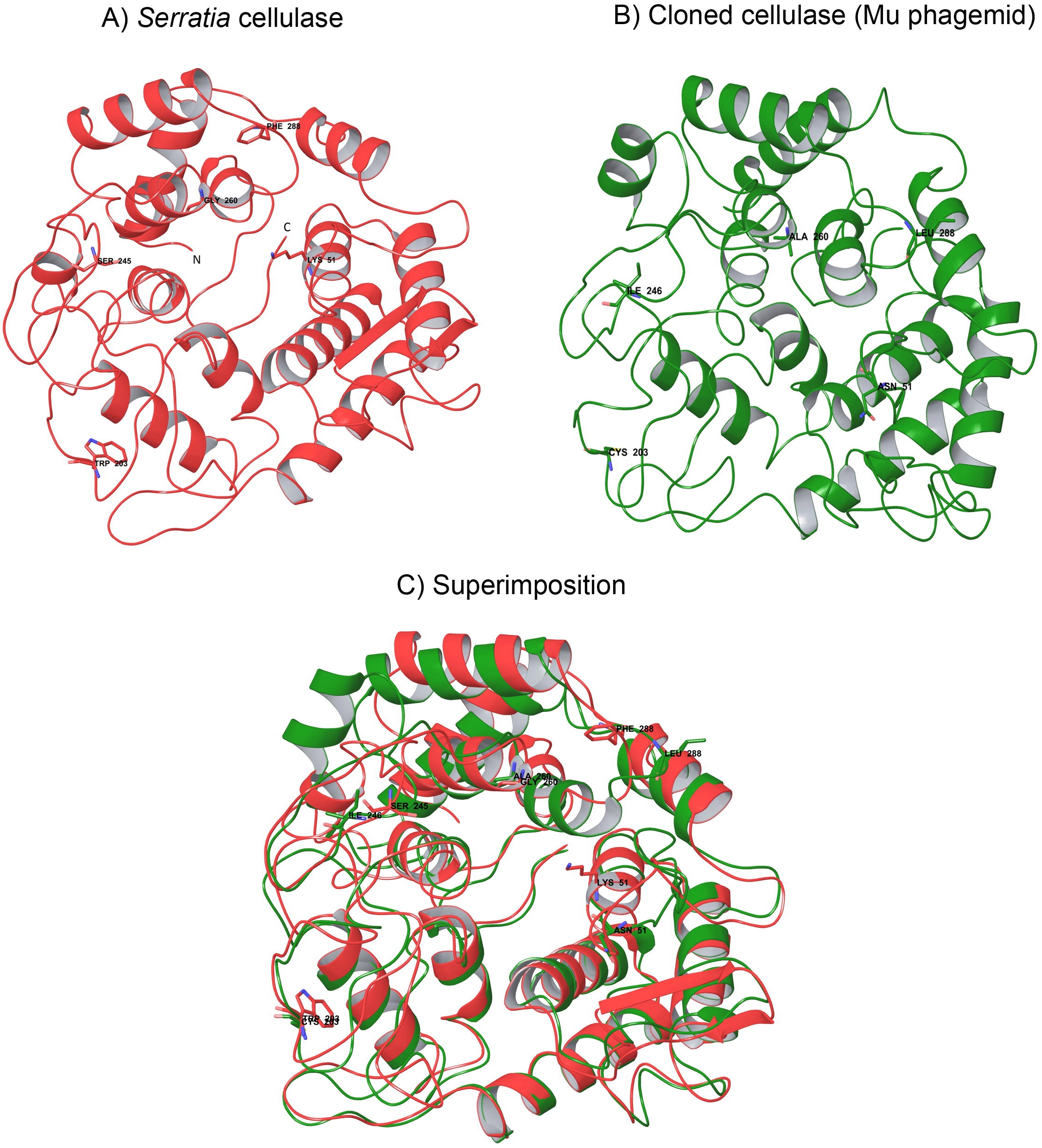
Three dimensional structures of Endoglucanase protein predicted by I-TASSER. A. Ribbon representation of endoglucanase in wildtype (*S. liquefaciens*). B. Cloned endoglucanase (Mu Phagemid). C. Superimposed structures of endoglucanase of wildtype (*S. liquefaciens*) and cloned endoglucanase (Mu Phagemid). The non-synonymous mutations were found at residues 51 (Lys - Asn), 203 (Trp-Cys), 246 (Thr-Iso), 260 (Gly-Ala) and 288 (Phe-Leu) are shown.

As of now, none of the crystal structure of cellulase of *Serratia* is available in PDB. Hence, a threading based model was predicted by I-TASSER. It generated the model using the structure of the bacterial cellulose synthase subunit, BcsZ (PDB: 3QXF) (Mazur and Zimmer 2011) as a template protein, which is a periplasmic protein with endo-β-1,4-glucanase activity and belongs to GH-8. The modeled endoglucanase protein of *Serratia* has 13 helices and two beta-sheets that are modified as loops in cloned cellulase protein though the amino acid sequences are exactly identical (Fig. 4A & B). The wildtype and cloned endoglucanase belongs to GH-8 that adopts (α/α)_6_ barrel folds. The predicted structures of wildtype endoglucanase has 83.5% of residues are in most favored regions and 14.7% are in allowed region. In case of cloned endoglucanase, 79% residues are in most favored regions and 16.9% are in the allowed region (Supplementary Figure 2). The structures of cloned and wildtype endoglucanases were superimposed and the root mean square deviation is 1.8Å (Fig. 4C).

There were differences observed in the molecular surface areas of the wildtype and cloned endoglucanase. The active site of wildtype endoglucanase is a narrow groove which lies parallel to the central axis whereas cloned endoglucanase structure is broad and tilted to ∼70° from the central axis (Fig. 5).

**Fig. 5.**
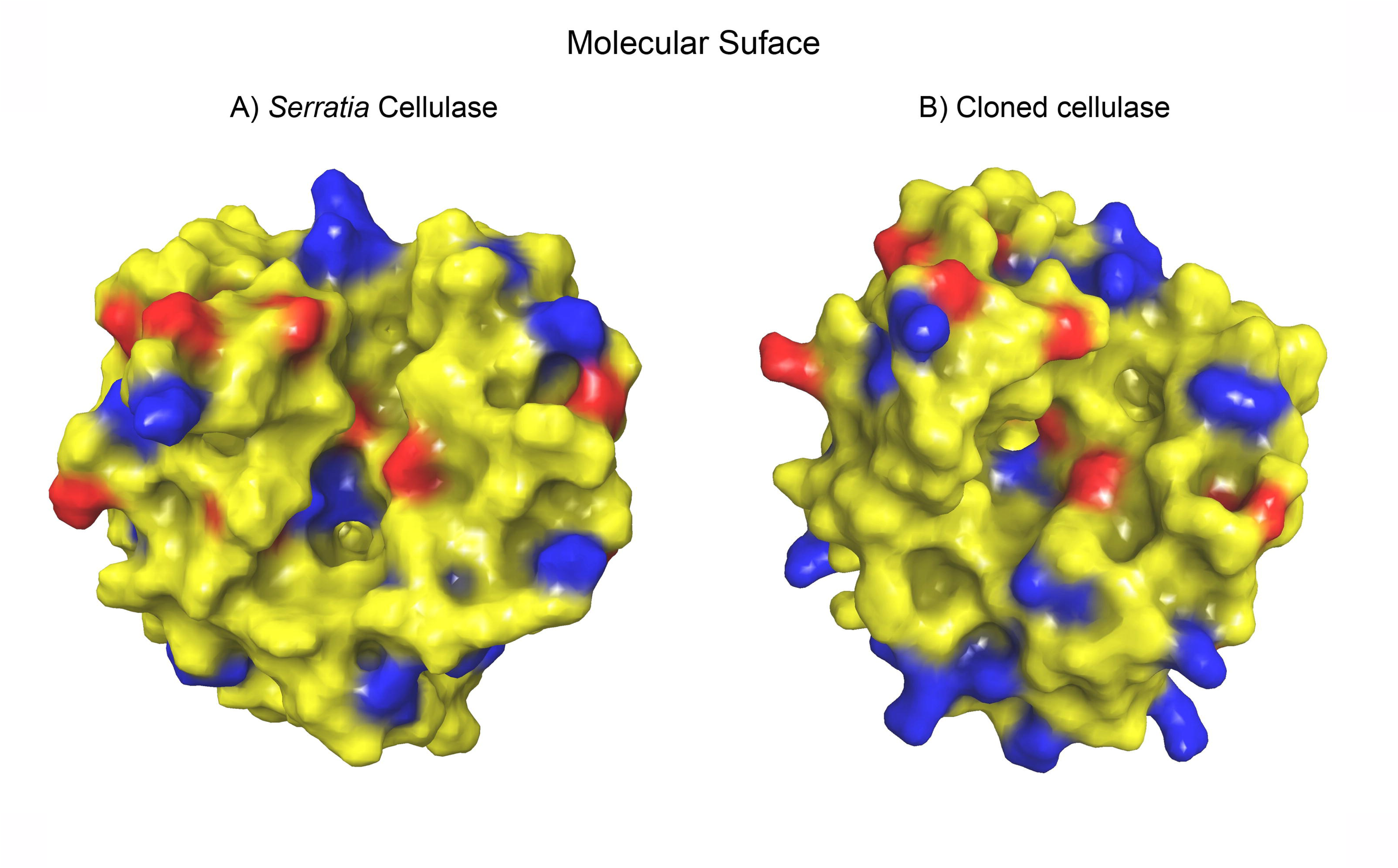
Solvent accessible surface of Endoglucanase proteins. A) Wildtype endoglucanase (*S. liquefaciens*). B) Cloned endoglucanase (Mu Phagemid). The surfaces are represented in residue charge property. The active sites are distinct in both the models and found tilted in cloned endoglucanase.

#### Discussion

Cellulose is the major component found in abundance in Earth, which could be harnessed and used as biofuels. Cellulase is the enzyme responsible for the hydrolysis of cellulose. Cellulolytic microorganism has the necessary enzymes to hydrolyze cellulose (Beguin 1990; Bayer et al. 1998; Lynd et al. 2002). There were only few reports available on *S. liquefaciens* as cellulolytic bacteria (Prem Anand and Sripathi 2004; Alwin Prem Anand et al. 2010; Hastuti et al. 2014). Due to higher cellulolytic activity, *S. liquefaciens* (Alwin Prem Anand et al. 2010) has been used in this experiment.

Endoglucanase are enzymes that hydrolyze cellulose by digesting β-1,4 glycosidic linkage (Lynd et al. 2002). The molecular mass of β-1,4 endoglucanase reported from our experiment is 32kDa, which is similar to the other β-1,4 endoglucanase from *Bacillus subtilis* (Lo et al. 1988), *Pectobacterium chrysanthemi* (Park et al. 2002), *Achlya ambisexualis* (Loprete and Hill 2002) and *Fomitopsis pinicola* (Yoon et al. 2008).

##### Sequence analysis of wildtype and cloned endoglucanase

Computational analyses have been made to unearth the rationale behind the increasing activity of cloned endoglucanase, as the sequence and structural composition of the protein determine its activity. The conserved domain of *S. liquefaciens* endoglucanase belongs to glycosyl hydrolyse family 8 (GH8, Fig. 2A). The GH8 enzymes hydrolyze β-1, 4 glycosidic bonds present in cellulose (Henrissat 1991; Henrissat and Bairoch 1993; Petersen et al. 2009). The nucleotide sequence analysis of endoglucanase of wildtype and the cloned gene revealed mutations at residues 51 (Lys - Asn), 203 (Trp-Cys), 246 (Thr-Iso), 260 (Gly-Ala) and 288 (Phe-Leu) (Fig. 2B). Among these mutations, except Gly260Ala mutation all other mutations are non-conservative.

In this study, we found increase in hydrophobic (Alanine, Isoleucine, Leucine) and polar (Asparagine, Cysteine) amino acids in cloned endoglucanase; but the additions were not in the catalytic domain. At the same time, reduced number of hydrophobic (Glycine, Phenylalanine), charged (Lysine) and polar (Threonine and Tryptophan) amino acids was observed in cloned endoglucanase (Table 2). It was said that protein stability is maintained by amino acid composition, hydrophobicities, compactness, polar and non-polar contributions to surface areas, hydrogen bonds in main- and side-chains, insertion/deletion of proline and salt bridges (Gromiha et al. 1999; Kumar et al. 2000). However, Sadeghi and colleagues (2006) reported that certain amino acids substitutions has no role in thermostability, but obtained through many minor structural modifications. The sequence analysis of both the proteins reveals the subtle differences in their molecular weight, theoretical pI, and hydropathicity (Table 2). Though there were only subtle changes, introduced by the changes in amino acid of the cloned endoglucanase; the stability and high enzyme activity at optimum temperature and pH is through structural modifications (described below).

DiANNA and CYSPRED prediction shows that Cys203 is bonded with Cys120. Thus, the connectivity of cysteine residues were altered between wildtype endoglucanase (Cys120 – Cys142, Cys141 – Cys235, Cys221 – Cys225) and cloned endoglucanase (Cys120 – Cys203, Cys142 – Cys221, Cys225 – Cys235). Moreover, mutation at 203^rd^ position introduces one more cysteine residue in the cloned endoglucanase in addition to existing 12 cysteine residues (Table 3). Based on the prediction, the cysteine substitution at 203^rd^ position in the cloned glucanase is responsible for the stability. Our results are in consistent with the study of Nemeth and colleagues (2002), who reported that the introduction of a disulfide bridge showed an increased conformational stability. This is corroborated by the temperature results where the cloned cellulase shows higher enzyme activity than wildtype at optimal temperature.

##### Structural analysis of cloned and wildtype endoglucanase

Several reports have demonstrated that the change in amino acid influenced the structural and functional properties of the endoglucanase enzymes. Amino acid residues His-131 and Glu-169 of endoglucanase were predicted to form the active site in the catalytic domain (Henrissat et al. 1995; Henrissat et al. 1996). When the amino acid at His-131 was mutated it results in the complete abolishment of enzyme activity, whereas mutation at Glu-169 resulted in significant loss of enzyme activity (Park et al. 1993). Henceforth, the theoretical models of wild and mutant type endoglucanase were generated to interrogate the structural consequences of amino acid substitutions in their secondary and tertiary structures. The secondary structures for both wildtype and mutated proteins were predicted using PSIPRED and found that the secondary structures of wildtype and cloned endoglucanase remain unaltered although five mutations were introduced in the sequence (Fig. 3A,B).

The 3D structure predicted by I-TASSER shows that wildtype and cloned endoglucanase belongs to GH-8 that adopts (α/α)_6_ barrel folds. The modeled endoglucanase protein of *Serratia* has 13 helices and two beta-sheets that are modified as loops in cloned cellulase protein though the amino acid sequences are exactly identical (Fig. 4A, B), while the structures of cloned and wildtype endoglucanases were superimposed and the root mean square deviation is 1.8Å (Fig. 4C). This shows that the structural models do not differ in their backbone, differ only in their side chains and its orientation. The Z-score of the pairwise structural alignment is 33.2 which means there is a significant structural similarity between the two structures. According to Yennamalli and colleagues (Yennamalli et al. 2011), thermostability in endoglucanase is fold-specific than protein families. Endoglucanase has three distinct structure folds: (α/β)_8_ fold, β-jelly roll fold and the (α/α)_6_ fold. The (α/α)_6_ fold has the substrate binding cleft ‘tunnel shaped’, which utilizes the ‘inverting mechanism’ for hydrolyzing β-1,4 glycosidic bonds, where the enzyme mechanism are not completely known. If we have to take into account, our endoglucanase belongs to GH8 with the (α/α)_6_ fold and hydrolyze cellulose via ‘inverting mechnaism’.

The stability of the enzymes depends on the maximum favourable interactions, forming a tightly packed hydrophobic core, exposing hydrophilic groups and optimizing intramolecular hydrogen bonding (Beadle and Shoichet 2002). Hence, both the predicted models are analyzed for number of hydrogen bonds and accessible surface area (ASA), as these parameters are crucial for functional properties of an enzyme. There is no difference in intramolecular hydrogen bonding in the predicted structures. But, the models differ in their ASA. The difference is mainly lies on the ASA of side chains than the ASA of backbone (Table 4). Few substitutions in the cloned endoglucanase have increased the total ASA that has enhanced the polar ASA and reduced the non-polar ASA. The molecular surface areas of cloned and wildtype endoglucanase were viewed and found that the active site of wildtype endoglucanase is a narrow groove which lies parallel to the central axis whereas cloned endoglucanase structure is broad and tilted to ∼70° from the central axis (Fig. 5).

**Table 4.** Structural analysis of the wildtype endoglucanase (S. liquefaciens) and cloned endoglucanase (Mu phagemid). Both the predicted models are analyzed for number of hydrogen bonds and surface accessible area. There is no difference in number of residues involved in hydrogen bonding. But the models differ in their accessible surface area (ASA). The difference is mainly lie on the ASA of sidechains than the ASA of backbone. Interatomic distances of selected residues in the active site of wildtype and cloned endoglucanase are found to be distinct.

| Parameters |  | *S. liquefaciens* | Mu Phagemid |
| --- | --- | --- | --- |
| **Secondary Structure** | Helix | 159 (50%) | 146 (46%) |
|  | Beta | 38 (12%) | 42 (13%) |
|  | Coil | 115 (36%) | 124 (39%) |
|  | Turn | 28 (8%) | 36 (11%) |
| **Hydrogen bonds (hbonds)** | Mean hbond distance | 2.3 sd=0.4 | 2.2 sd=0.4 |
|  | Mean hbond energy | -1.5 sd=1.1 | -1.4 sd=1.1 |
|  | Residues with hbonds | 232 (74%) | 233 (74%) |
| **Accessible surface area (ASA)** | Total ASA | 15037.5 Å<sup>2</sup> | 15121.8 Å<sup>2</sup> |
| **Inter atomic distances (Å)** | Cys110-Pro307 | 16.45 | 15.52 |
|  | Lys51-Cys221 | 14.60 | 18.74 |
|  | Pro289-Val259 | 8.89 | 13.36 |
|  | Lys140-Tyr143 | 8.37 | 7.87 |

Exo and endoglucanases differ only in their binding site for endoglucanase, which is formed of loops. In case of exonucleases, a tunnel like structure encloses the catalytic residues whereas in endoglucanases, the binding cleft, formed of shorter loops, has direct access to cellulose chains (Ubhayasekera et al. 2005; Juturu and Wu 2014). The wildtype and cloned endoglucanase belongs to GH8, which has (α/α)_6_ barrel architecture similar to endoglucanase CelA of *Clostridium thermocellum* (Alzari et al. 1996; Guerin et al. 2002), BcsZ (PDB: 3QXF) (Mazur and Zimmer 2011) and endoglucanase Cel10 from *K. pneumoniae* (Attigani et al. 2016). The active site of GH8 enzyme is a cleft, which allows substrate binding and cleavage (Henrissat 1991; Henrissat and Bairoch 1993; Petersen et al. 2009). The active site was manually inspected and compared between the wildtype and cloned endoglucanases as there are no much reports on endoglucanase of *S. liquefaciens*. The catalytic domain (CD) (Fig. 6) and the inter-atomic distances have been measured to show the difference in the active sites of wildtype and cloned endoglucanase (Table 4). Though the mutated amino acids are not in the active sites, it has altered the cleft of cloned endoglucanase compared to the wildtype endoglucanase. In cloned endoglucanase, the cleft is wide and the inter-atomic distances were found to be longer. It is known that open or extended cleft is usually found in amorphous cellulose degrading enzymes, such as endoglucanase (Lynd et al. 2002; Boraston et al. 2004; Payne et al. 2015). The enzyme-substrate binding site in the BcsZ (Mazur and Zimmer 2011), endoglucanase CelA (Alzari et al. 1996; Guerin et al. 2002) and endoglucanase Cel10 (Attigani et al. 2016) is similar to that of our wildtype and cloned endoglucanase pointing out that these are open substrate-binding cleft. The widening of the cleft in the cloned endoglucanase favours increased enzyme activity.

**Fig. 6.**
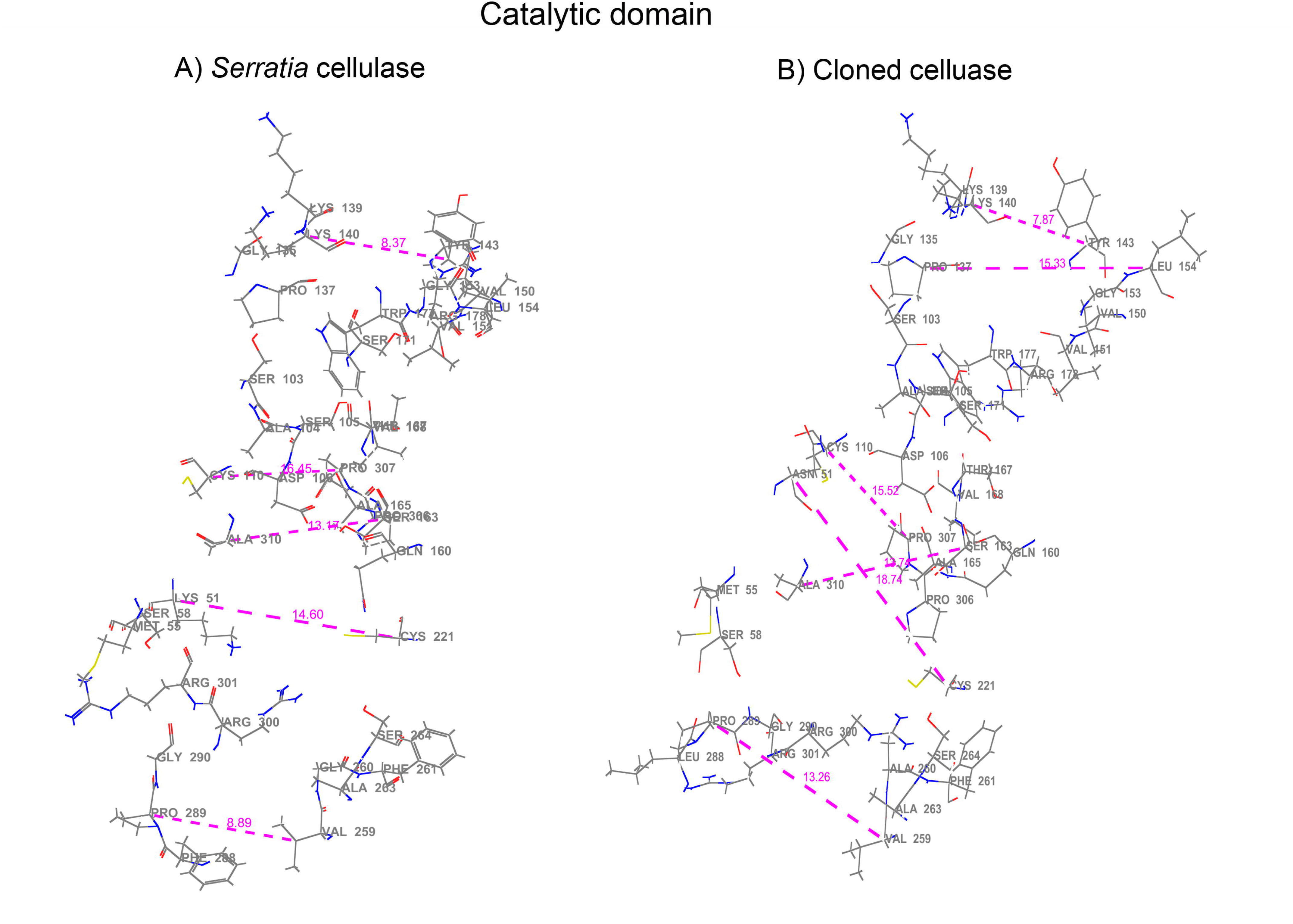
The active site of A) wildtype endoglucanase (*S. liquefaciens*) and B) cloned endoglucanase (Mu phagemid). The interatomic distances of amino acids lying in the right and left side of the active sites are displayed and found to be altered in cloned endoglucanase.

The cloned enzyme have shown 24% increased activity compared to wildtype endoglucanase, bioinformatics analysis of wild and mutant endoglucanase did not show dramatic differences in their molecular models. There was no significant difference observed in their molecular weight, pI, hydropathy scale and secondary structural elements. However, one of the substitutions have introduced one cysteine residue, which was found to be involved in altered disulfide bridge formation based on *in silico* prediction. The active sites of both enzymes was found to be altered especially differ in their interatomic distances, which might have significant role in interactions with substrates.

In conclusion, the computational analysis of wildtype and cloned endoglucanase proteins showed that the latter changed in its amino acid composition, iso electric point and number of cysteine residues. The structures also revealed the subtle variations in the composition of secondary structures, accessible surface area, and interatomic distances in the active sites, which are crucial structural features responsible for increased enzyme activity of endoglucanase. We conclude the reason for increased enzyme activity in the cloned endoglucanase might be due to the structural orientation conferred by the amino acid changes and widening of the cleft favours increased enzyme activity. However, only the predicted models have been used in this work, further molecular dynamic study is important to provide more insights. To conclude, the structural consequences were the reason for high enzymatic activity in the cloned enzyme. The attempt made in the present study would pave a way for producing more flexible and, enzyme with high stability.

## Acknowledgement

The authors thank Prof. Munavar, Madurai Kamaraj University for providing the necessary microbial strains for the research.

## Author contribution

GS conducted the experiments, analyzed the data, produced figures, draft the manuscript and critically revised the manuscript. AMJ performed the *in silico* analysis, analyzed the data, produced figures and incorporated the *in silico* results. SJV participated in the experimental design, analyzed the data and helped to draft the manuscript. APA conceived the study, designed the experiments, analyzed the data, draft the manuscript and critically revised the manuscript. All author approve the final manuscript for publication.

## Supporting information legend

**Supplementary Figure 1. Phylogenetic analysis of cellulase proteins of *S. liquefaciens* with Enterobacteriaceae family.**

The tree depicts the phylogenetic relationship of *S. liquefaciens* with other species of *Serratia* and Enterobacteriaceae family.

**Supplementary Figure 2. Ramachandran plot obtained for predicted endoglucanase protein in PROCHECK.**

A) Endoglucanase of wildtype endoglucanase (*S. liquefaciens*). 83.5% of residues are in most favoured regions and 14.7% are in allowed region. B) Ramachandran Plot for cloned endoglucanase (Mu Phagemid). Out of 312 amino acids, 211 residues are in most favoured regions and 16.9% are in allowed region. Glycines are marked as triangles. The most favoured regions, allowed regions are in red and yellow respectively.

