## Supplementary figures and images for "*In vivo* cloning of β 1-4 endoglucanase gene of *Serratia liquefaciens* using Muduction and its *in silico* analysis"

### Supplementary Materials

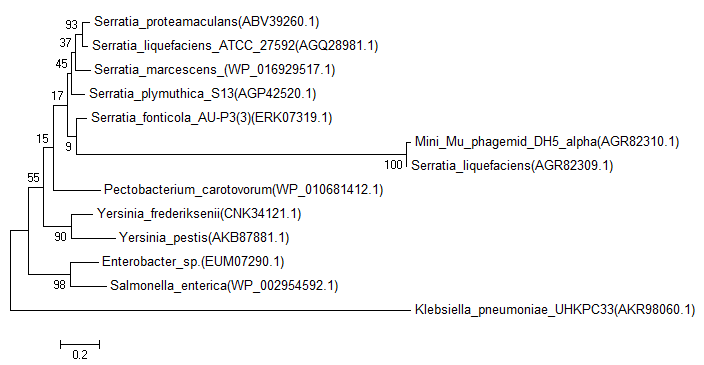

### Supplementary Materials

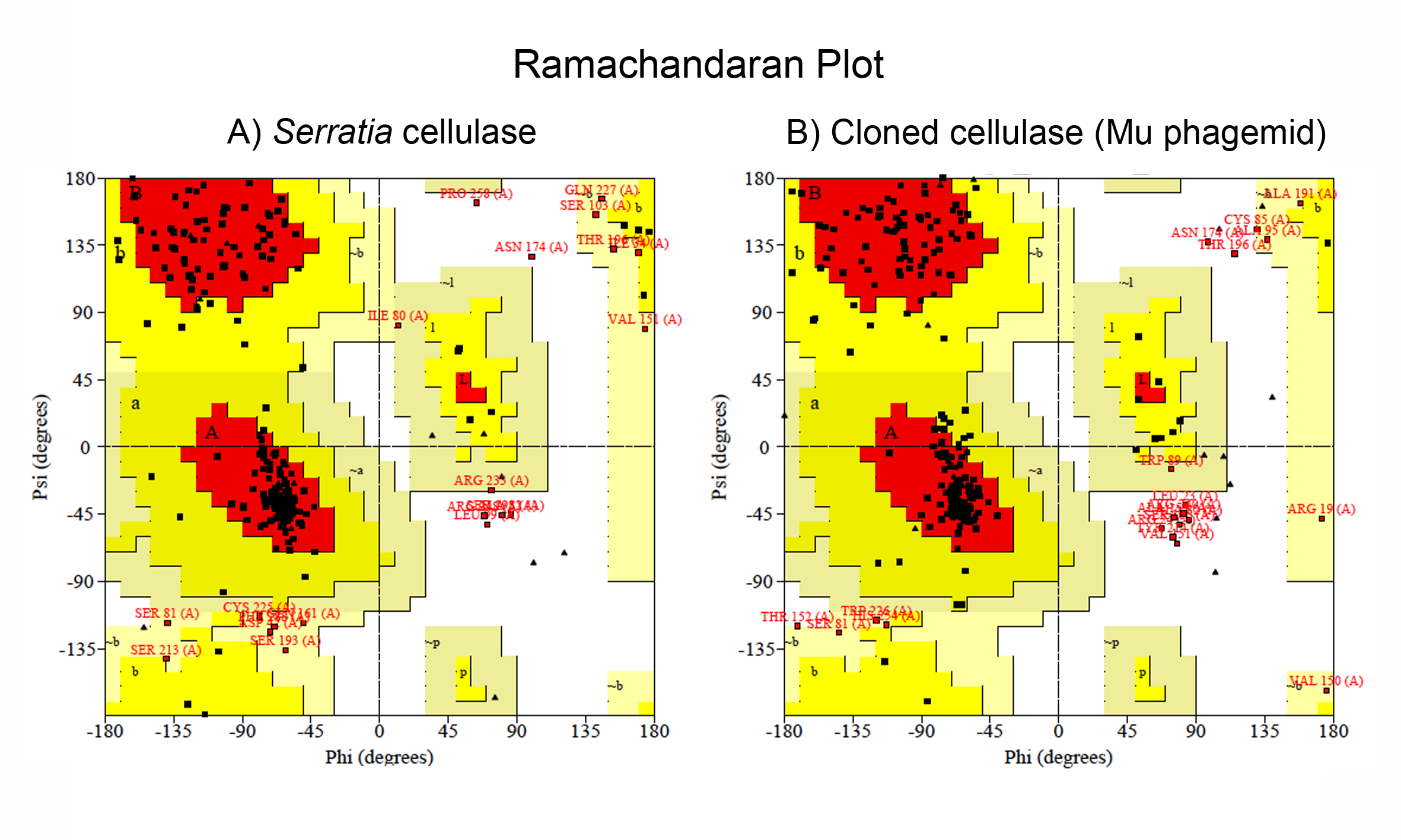
